# Enhancement of the immunogenicity of a *Mycobacterium tuberculosis* fusion protein using ISCOMATRIX and PLUSCOM nano-adjuvants after nasal administration in mice

**DOI:** 10.1101/2021.08.27.458002

**Authors:** Arshid Yousefi Avarvand, Zahra Meshkat, Farzad Khademi, Ehsan Aryan, Mojtaba Sankian, Mohsen Tafaghodi

**Affiliations:** Department of Laboratory Sciences, School of Allied Medical Sciences, Ahvaz Jundishapur University of Medical Sciences, Ahvaz, Iran; Antimicrobial Resistance Research Center, Mashhad University of Medical Sciences, Mashhad, Iran; Department of Medical Bacteriology and Virology, School of Medicine, Mashhad University of Medical Sciences, Mashhad, Iran; Department of Microbiology, School of Medicine, Ardabil University of Medical Sciences, Ardabil, Iran; Immunobiochemistry laboratory, Immunology Research Center, Bu-Ali Research Institute, Mashhad, Iran; Nanotechnology Research Center, Pharmaceutical Technology Institute, Mashhad University of Medical Sciences, Mashhad Iran

**Author notes:** Corresponding author Official email address for corresponding author (Mohsen Tafaghodi). Zahra Meshkat official.

**Keywords:** *Mycobacterium tuberculosis*, HspX/EsxS, ISCOMATRIX, PLUSCOM, MPLA, Nasal administration

## Abstract

**Background:** Tuberculosis (TB), a contagious disease caused by *Mycobacterium tuberculosis* (*M. tuberculosis*), remains a health problem worldwide and this infection has the highest mortality rate among bacterial infections. Current studies suggest that intranasal administration of new tuberculosis vaccines could enhance the immunogenicity of *M. tuberculosis* antigens. Hence, we aim to evaluate the protective efficacy and immunogenicity of HspX/EsxS fusion protein of *M. tuberculosis* along with ISCOMATRIX and PLUSCOM nano-adjuvants and MPLA through the intranasal administration in mice model.

**Methods:** In present study, the recombinant fusion protein was expressed in *Escherichia coli* and purified and used to prepare different nanoparticle formulations in combination with ISCOMATRIX and PLUSCOM nano-adjuvants and MPLA. Mice were intranasally vaccinated with each formulation three times at an interval of 2 weeks. Finally, IFN-γ, IL-4. IL-17 and TGF-β concentration in supernatant of cultured splenocytes of vaccinated mice as well as serum titers of IgG1 and IgG2a and sIgA titers in nasal lavage were determined.

**Results:** According to obtained results, intranasally vaccinated mice with formulations containing ISCOMATRIX and PLUSCOM nano-adjuvants and MPLA could effectively induced IFN-γ and sIgA responses. Moreover, both HspX/EsxS/ISCOMATRIX/MPLA and HspX/EsxS/PLUSCOM/MPLA and their BCG booster formulation could strongly stimulate the immune system and enhance the immunogenicity of *M. tuberculosis* antigens.

**Conclusion:** The results demonstrate the potential of HspX/EsxS-fused protein in combination with ISCOMATRIX, PLUSCOM and MPLA after nasal administration in enhancing immune response against of *M. tuberculosis* antigens. So, nasal immunization with these formulations, could induce immune responses and considered as new TB vaccine or as BCG booster.

## Introduction

Tuberculosis (TB) is a contagious disease with approximately 1.7 billion latently infected people and over 1.2 million deaths annually. TB is among the 10 causes of death worldwide, according to the latest World Health Organization (WHO) report which can be controlled using early vaccination as well as rapid detection and treatment with the first- and second anti-TB drugs (1-4). However, the emergence of *Mycobacterium tuberculosis* (*M. tuberculosis*) resistant strains particularly rifampicin-resistant and multidrug-resistant TB (MDR) have led to treatment failures (5). Furthermore, for many years, existence of some disadvantages in the only licensed *M. tuberculosis* vaccine, BCG (Calmette-Guérin Bacillus), has led to many efforts to assess the other ways of controlling the TB disease (6, 7). Efficacy of BCG vaccine for pulmonary TB decreases during lifetime and therefore it is more effective against newborns and children (8, 9). Additionally, BCG is not recommended for patients with immune deficiency and is not able to control the latent TB infection which can act as reservoir of active TB infection (3). Therefore, several vaccines are in different steps of clinical or preclinical studies. These vaccines are examined for either pre-exposure prevention which can be administrated before TB infection in newborns and adolescents or as post-exposure and therapeutic vaccines which can be administered in adolescents and adults after TB infection to eliminate latent TB. These new types of TB vaccines considered as either alternative for BCG vaccine or as booster of BCG prime (10-12). Multi-stage subunit vaccines as pre-exposure, post-exposure and therapeutic vaccines, are promising candidates for boosting BCG-primed immunity or a prime-vaccine alternative for BCG vaccine (10, 13). On the other hand, combining multi-stage subunit vaccines with adjuvants and delivery systems can potentiate the immunogenicity of multi-stage vaccines, protect antigens from enzymatic degradation and *in vivo* elimination, targeted delivery and then efficient uptake of antigens and control of antigens release (14, 15). In a series of the studies, we evaluated the potential of a novel multicomponent subunit vaccine candidate called HspX/EsxS-fused protein, a latent-phase protein (HspX) plus an early-phase protein (EsxS), along with various adjuvants such as DOTAP (1, 2-dioleoyl-3-trimethylammonium propane), MPLA (monophosphoryl lipid A) and DDA (dimethyl dioctadecylammonium bromide) as well as delivery systems such as PLGA (poly (lactide-co-glycolide)) through different administration routes in animal model in order to enhance the immunogenicity of *M. tuberculosis* antigens (16-18). Furthermore, two nano-adjuvants ISCOMATRIX, a negatively charged particle, and PLUSCOM, a positively charged ISCOMATRIX, were also evaluated along with HspX/EsxS-fused protein via subcutaneous administration and the results were promising in animal model (unpublished data). However, as *M. tuberculosis* enters via respiratory tract, the mucosal administration of these formulations might rapidly induce the innate and adaptive immune response at the at the respiratory mucosal surfaces (10, 19, 20). Therefore, we followed two aims; 1) determine the potential of HspX/EsxS-fused protein in combination with ISCOMATRIX, PLUSCOM and MPLA after nasal administration and 2) comparison of the current results with our previous results.

## Materials and Methods

### Preparation of HspX/EsxS protein, ISCOMATRIX and PLUSCOM nano-adjuvants

Synthesis of the HspX/EsxS fused protein was performed as described previously. To perform this, the recombinant fusion protein was expressed in *Escherichia coli*, purified on a chromatography column (Parstous Biotechnology, Iran) and then verified by SDS-PAGE and western blot. The protein concentration was also measured by BCA kit (Parstous Biotechnology, Iran) (16). Furthermore, ISCOMATRIX and PLUSCOM nano-adjuvants were prepared by the lipid film hydration method. Briefly, to provide the ISCOMATRIX nano-adjuvant, 200 µL of cholesterol (4 mg/mL) along with 320 µL of phosphatidylcholine (8 mg/mL) (Avanti polar lipids, USA) were dissolved in dichloromethane and then mixed and vacuum dried to eliminate the dichloromethane and establish the lipid film. The PLUSCOM lipid film was also prepared by mixing 200 µL of DDA (4, 8 or 16 mg/mL) and 320 µL of phosphatidylcholine (8 mg/mL) dissolved in dichloromethane. Both ISCOMATRIX and PLUSCOM lipid films were hydrated by 200 mg of sucrose (Merck, Germany), dissolved in distilled water (2 mL) and butanol (2 mL), and then freeze-dried for overnight. The freeze-dried powders were combined with an aqueous phase containing saponin (8 mg in 4 mL of PBS (0.01 M), pH 7.4) (Sigma-Aldrich, USA) and then bath sonicated (Kerry, UK) at 37 °C for 10 minutes. Dynamic light scattering (DLS) (Zetasizer Nano, Malvern, UK) was used to measure the particle size and surface charge of nano-adjuvants (20-23).

### Preparation of vaccine formulations, mice vaccination and immunoassay

The following vaccine formulations were prepared in aseptic conditions in order to nasal administration of 10 mice groups including 5 BALB/c mice, 6 to 8 weeks old, in each group: 1) PBS (negative control), 2) BCG (5×10^5^ CFU/mouse), 3) HspX/EsxS, 4) HspX/EsxS/MPLA, 5) HspX/EsxS/ISCOMATRIX, 6) HspX/EsxS/PLUSCOM, 7) HspX/EsxS/ISCOMATRIX/MPLA, 8) HspX/EsxS/PLUSCOM/MPLA, 9) HspX/EsxS/ISCOMATRIX/MPLA as BCG booster and 10) HspX/EsxS/PLUSCOM /MPLA as BCG booster. Mice were nasally vaccinated with 20 µL of each formulation (10 µg of HspX/EsxS, 15 µg of ISCOMATRIX, 15 µg of PLUSCOM and 15 µg of MPLA) three times at an interval of 2 weeks.

Three weeks after final vaccination, nasal lavage, blood and spleen of vaccinated mice were used to assay IgA, IgG1 and IgG2a titers as well as interferon gamma (IFN-γ), interleukin 4 (IL-4), interleukin 17 (IL-17) and transforming growth factor beta (TGF-β) cytokines (18, 24). For cytokine assays, production of IFN-γ, IL-4, IL-17 and TGF-β by splenic lymphocytes (2 × 10^6^ cells/well) of mice which were stimulated with each formulation, measured in supernatant of cultured splenocytes according to the enzyme-linked immunosorbent assay (ELISA) kit (eBioscience, USA). Additionally, goat anti-mouse IgA:HRP, IgG1:HRP, and IgG2a:HRP (Invitrogen, USA) were used for measurement of lavage anti-HspX/EsxS IgA titers and serum anti-HspX/EsxS IgG1 and IgG2a titers(20).

## Statistical analysis

All statistical analysis was performed by the GraphPad Prism 8.0 software, and all data analysis was performed by one-way ANOVA in combination with Tukey’s multiple comparison tests. Values were expressed as mean ± SD, when p-value<0.05, differences was considered as statistically significant. Significance was presented as ^*^(P < 0.05), ^**^(P < 0.01), ^***^(P < 0.001) and ^****^(P < 0.0001) and not significant was shown as (ns).

## Results

### Assessment of IFN-γ response

After nasal administration, our results showed that formulations contain nano-adjuvants ISCOMATRIX (ISCOMATRIX/HspX/EsxS) and PLUSCOM (PLUSCOM/HspX/EsxS) were able to boost HspX/EsxS immunogenicity and induced higher level of IFN-γ response compared to HspX/EsxS alone, (P<0.001) and (P<0.0001) respectively. Also, addition of MPLA adjuvant to ISCOMATRIX/HspX/EsxS and PLUSCOM/HspX/EsxS formulations was promoted the immune responses. The spleen cells of the mice receiving HspX/EsxS/ISCOMATRIX/MPLA and HspX/EsxS/PLUSCOM/MPLA formulations were significantly produced the higher level of IFN-γ than those receiving HspX/EsxS/ISCOMATRIX and HspX/EsxS/PLUSCOM, respectively (p<0.01) and (p<0.05). There was no significant difference between BCG boosters of HspX/EsxS/ISCOMATRIX/MPLA and HspX/EsxS/PLUSCOM/MPLA (P >0.05), although, both HspX/EsxS/ISCOMATRIX/MPLA and HspX/EsxS/PLUSCOM/MPLA and their BCG booster formulation were able to induce IFN-γ response significantly higher than BCG group (P <0.001) (Figure 1).

**Figure 1.**
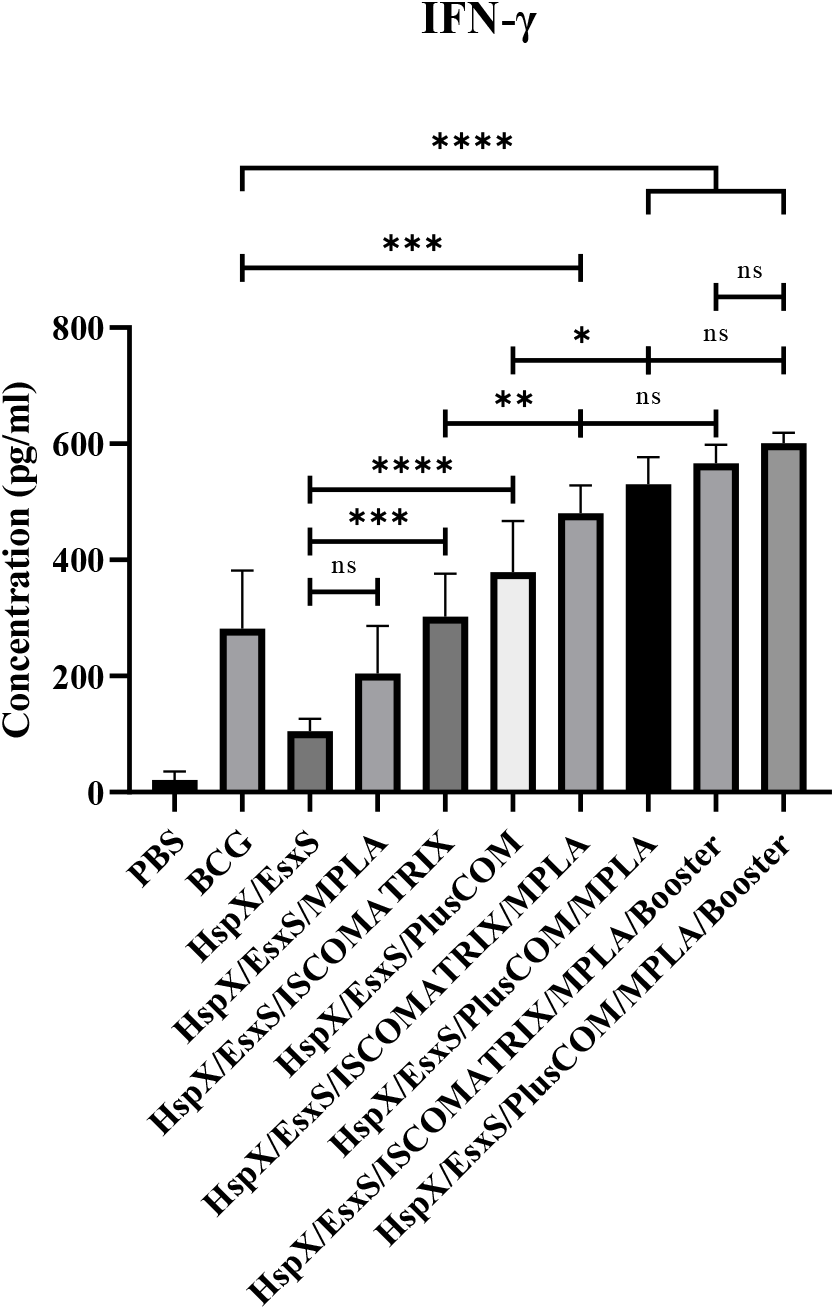
The level of IFN-γ produced in the spleen cells of the mice receiving different formulations.

### Assessment of IL-17 response

Our result show that different formulations did not induce IL-17 response significantly in the stimulated splenic lymphocytes of mice compare to BCG group (P >0.05) (Figure 2).

**Figure 2.**
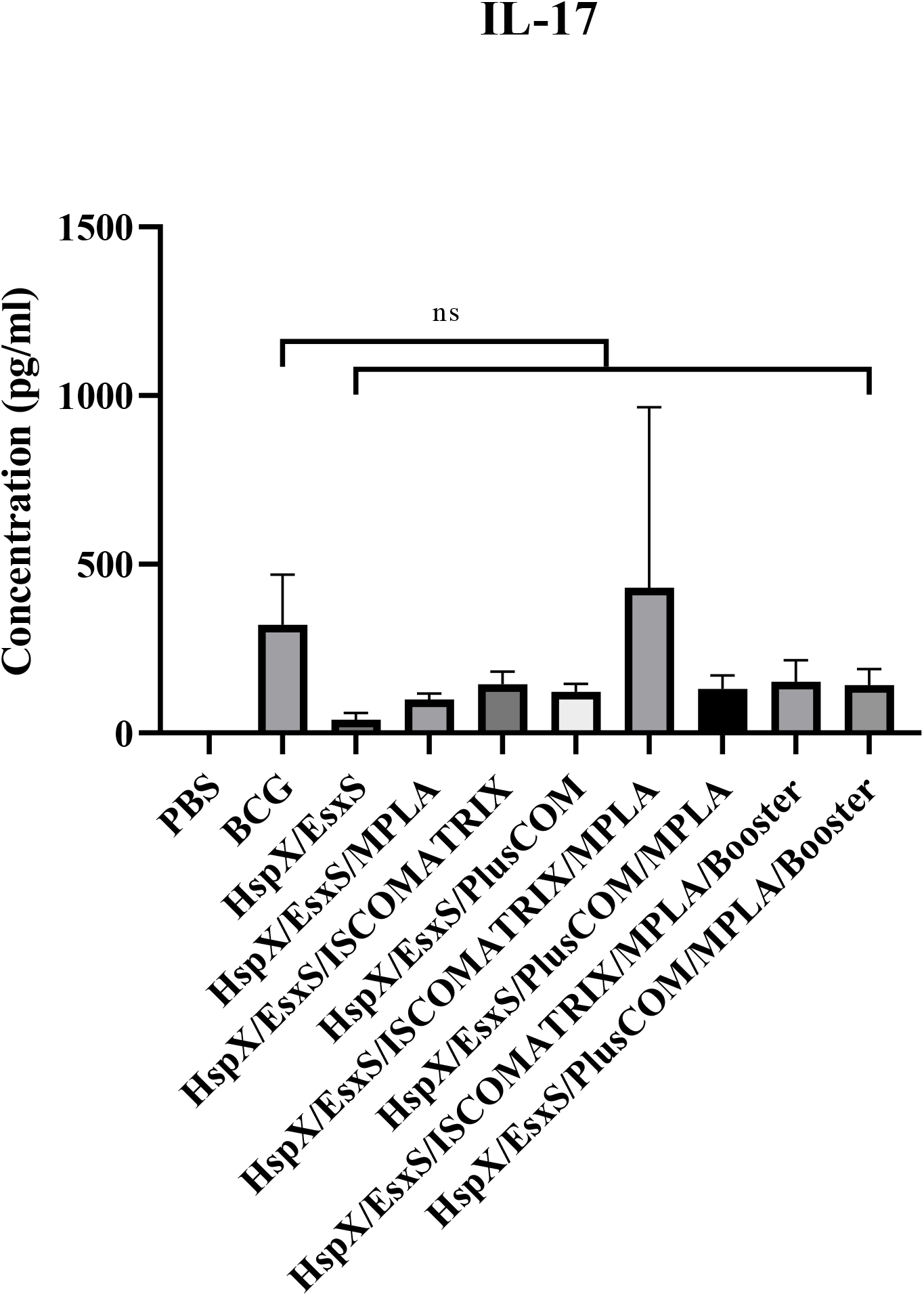
The level of IL-17 produced in the spleen cells of the mice receiving different formulations.

### Assessment of IL-4 response

According to obtained result the level of IL-4 secretion in vaccinated mice with BCG booster of HspX/EsxS/PLUSCOM/MPLA formulation was higher than HspX/EsxS and BCG vaccine (P>0.05). However, any formulations weren’t able to indue IL-4 response significantly higher than BCG group (P>0.05). (Figure 3)

**Figure 3.**
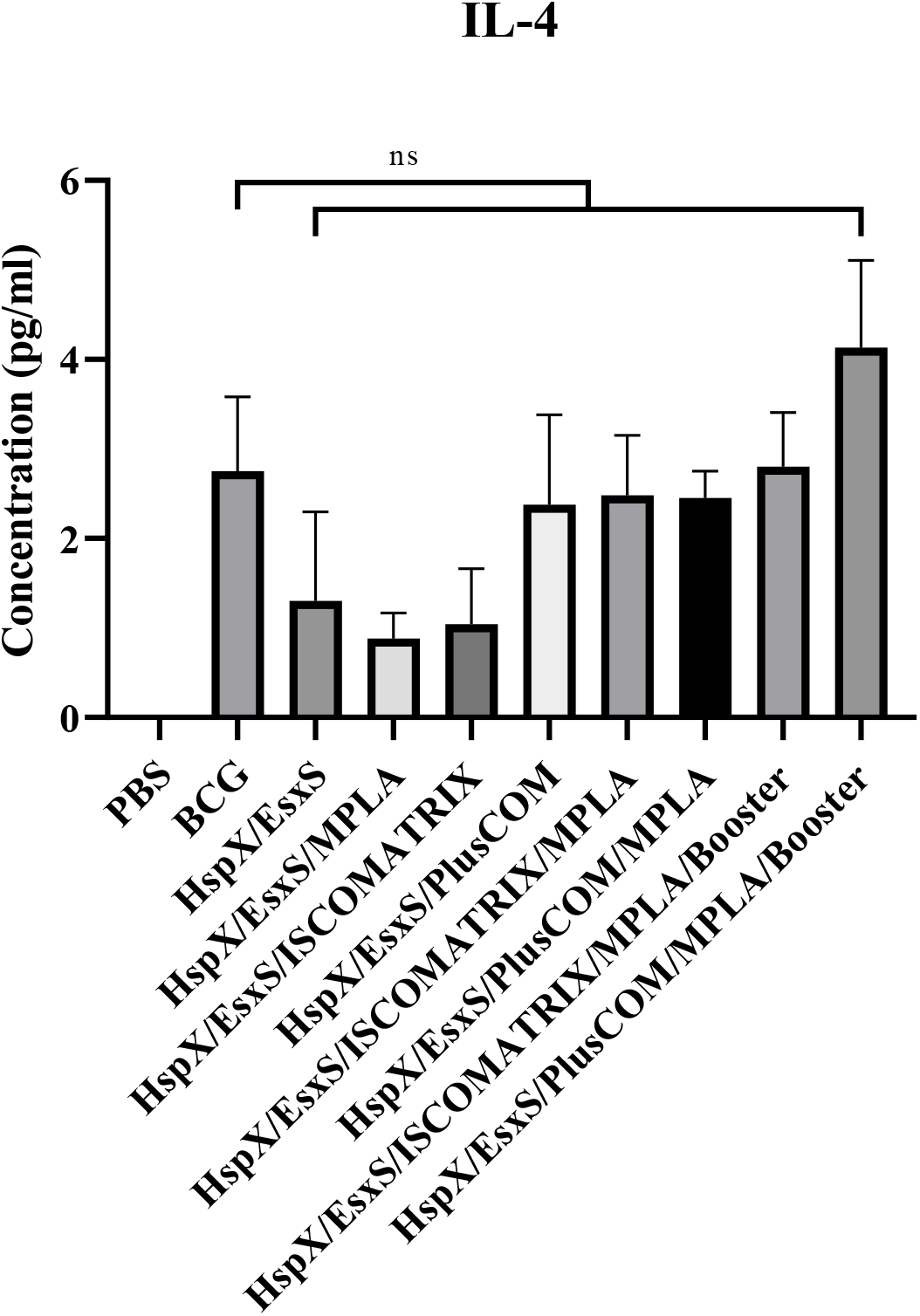
The level of IL-4 produced in the spleen cells of the mice receiving different formulations.

### Assessment of TGF-β response

Similar to IL-4 and IL-17, there was no significant difference between different formulation and BCG group in induction of TGF-β response (P>0.05). (Figure 4)

**Figure 4.**
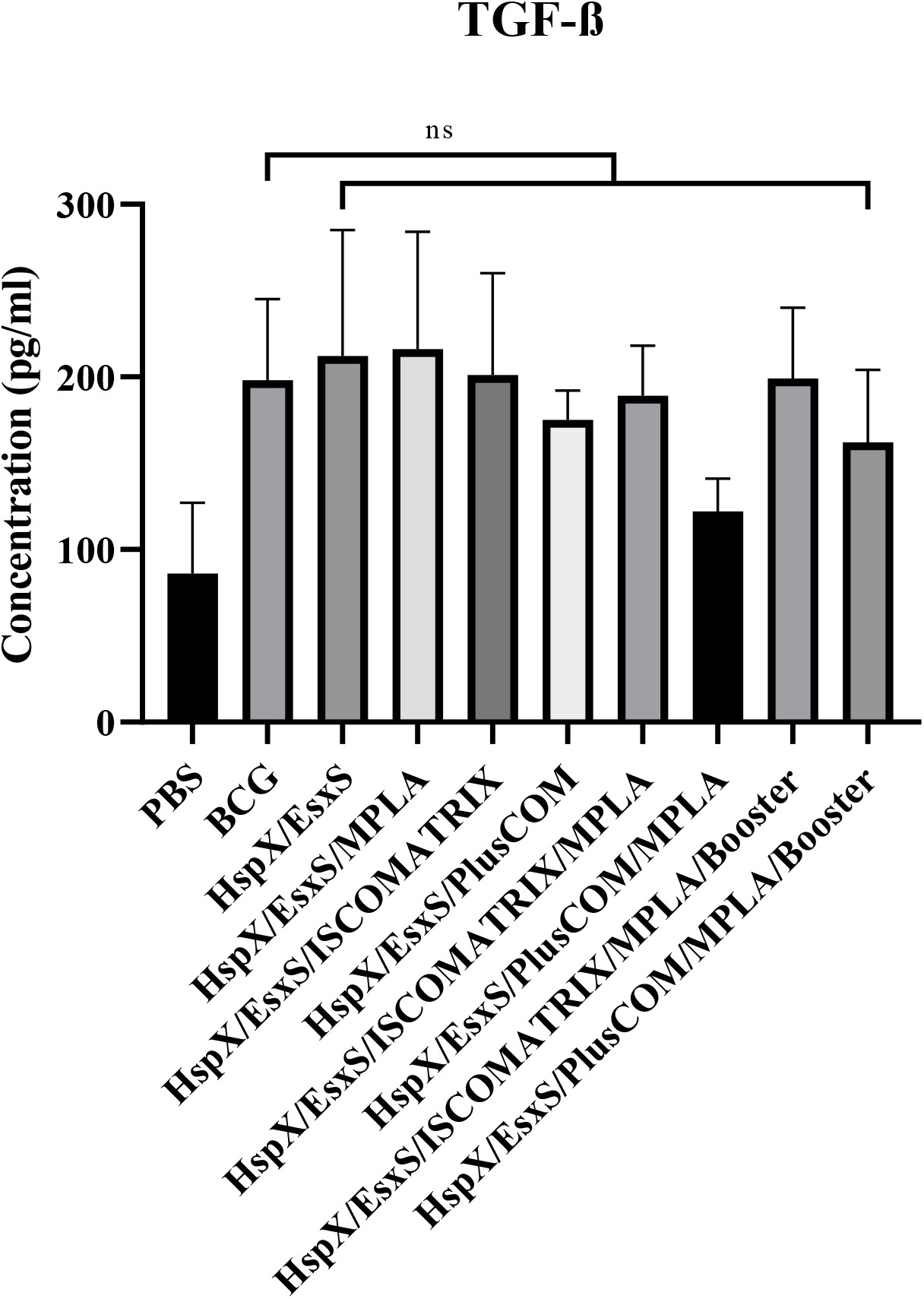
The level of TGF-β produced in the spleen cells of the mice receiving different formulations.

### Assessment of IgG2a antibody response

the level of serum anti-HspX/EsxS IgG2a titers in mice vaccinated with HspX/EsxS/ISCOMATRIX/MPLA, HspX/EsxS/PLUSCOM/MPLA and their BCG booster formulations was significantly higher than HspX/EsxS and BCG vaccine (P <0.0001). Additionally, HspX/EsxS/PlusCOM/MPLA/Booster was able to significantly increase IgG2a responses higher than HspX/EsxS/PlusCOM/MPLA (P <001). (Figure 5)

**Figure 5.**
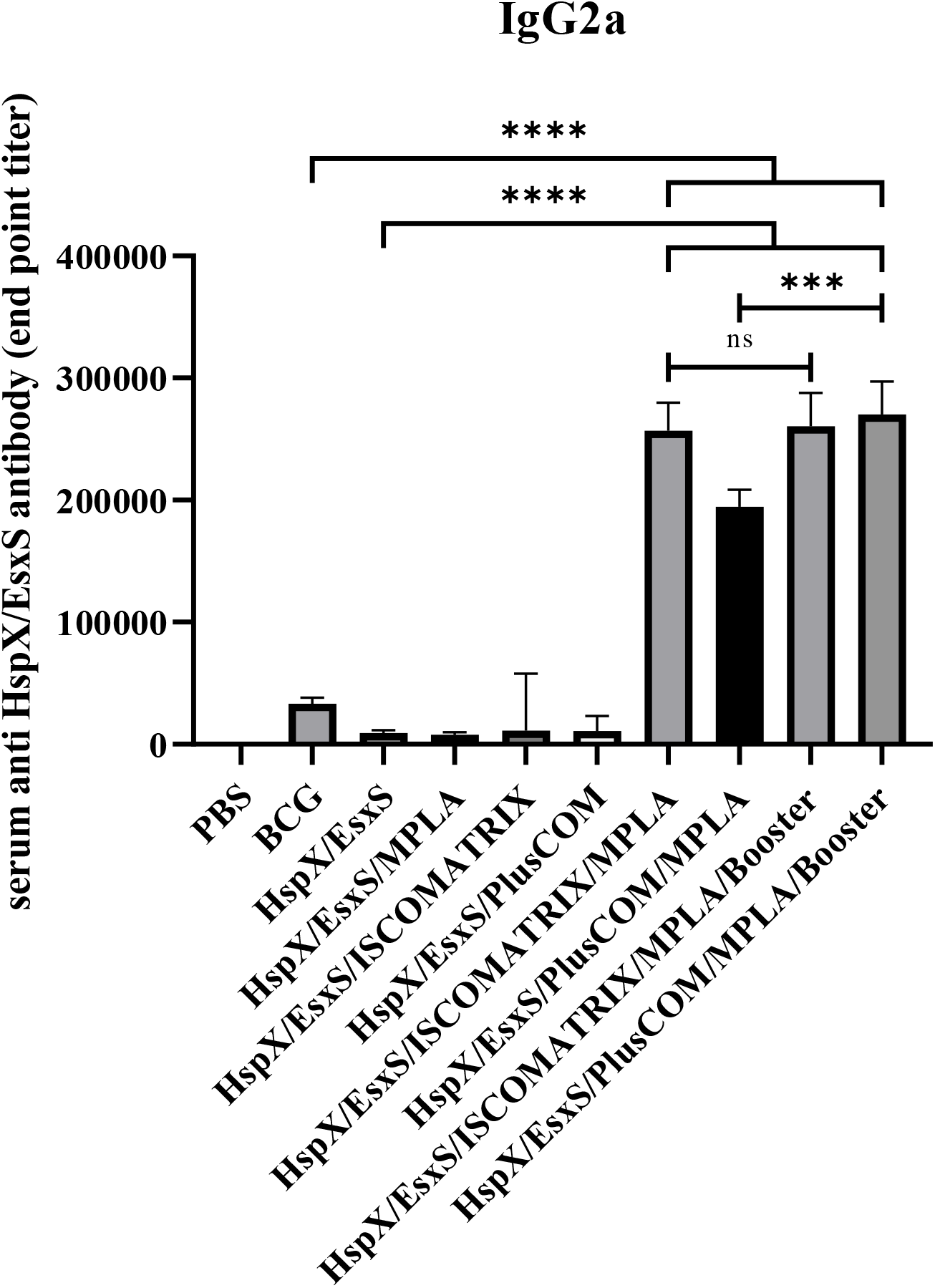
The level of IgG2a produced in the serum of mice receiving different formulation.

### Assessment of IgG1 antibody response

The level of serum anti-HspX/EsxS IgG1 titers as well as IgG2a was significantly increased in the mice receiving HspX/EsxS/ISCOMATRIX/MPLA, HspX/EsxS/PLUSCOM/MPLA and BCG booster formulations in comparison with HspX/EsxS and BCG vaccine (P <0.0001). Also, addition of MPLA adjuvant and BCG booster formulation, significantly increased the effect of HspX/EsxS/ISCOMATRIX and HspX/EsxS/PLUSCOM formulations on IgG1 antibody response (P <0.0001) (Figure 6).

**Figure 6.**
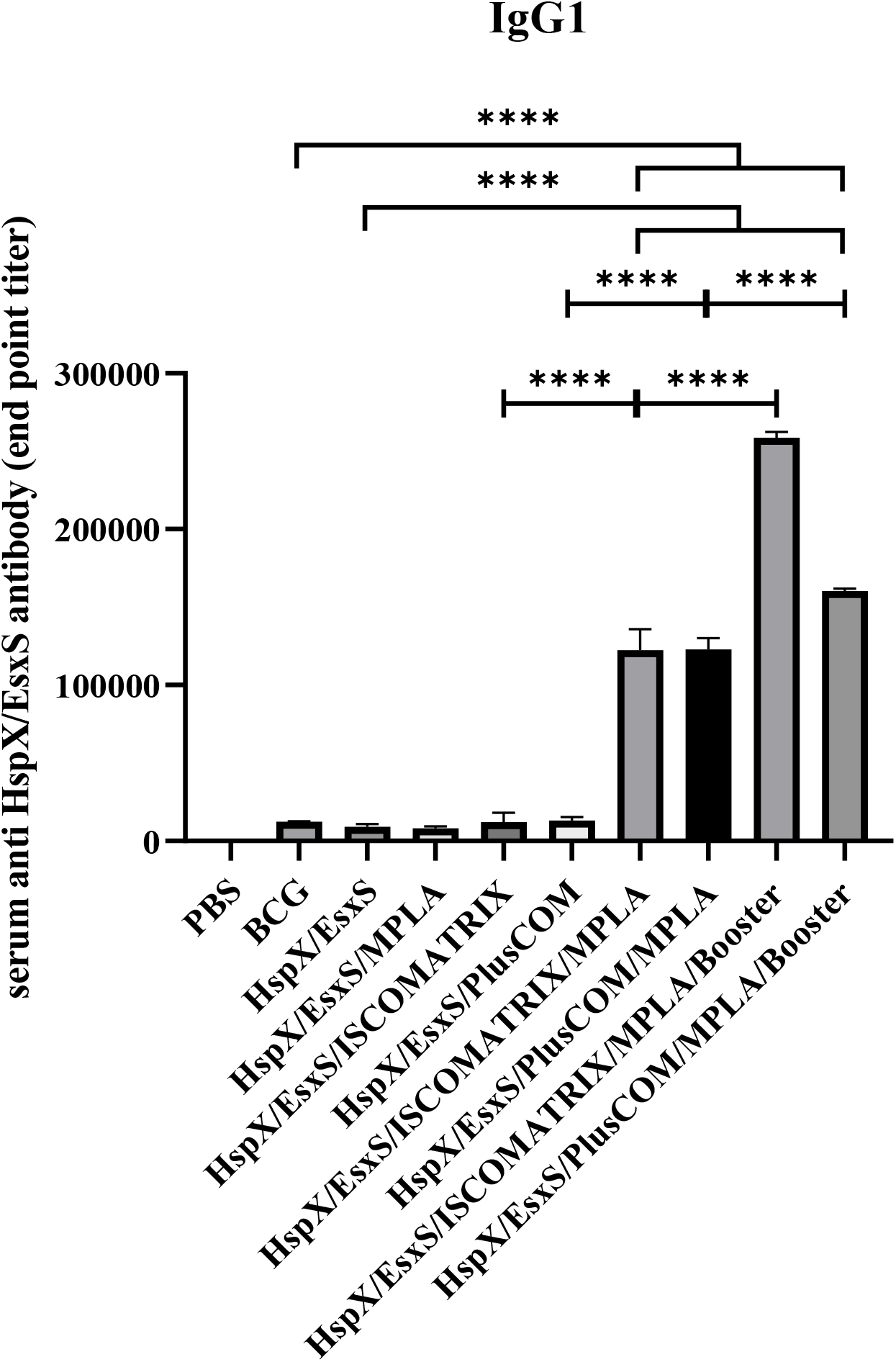
The level of IgG1 produced in the serum of mice receiving different formulation.

### Assessment of sIgA antibody response

Anti-HspX/EsxS sIgA antibody in nasal lavage of vaccinated mice was significantly higher in HspX/EsxS/ISCOMATRIX, HspX/EsxS/PLUSCOM, HspX/EsxS/ISCOMATRIX/MPLA, HspX/EsxS/PLUSCOM/MPLA and their BCG booster formulation in comparison with HspX/EsxS and BCG vaccine (P <0.05). Furthermore, the highest level of sIgA antibody response belonged to HspX/EsxS/PlusCOM/MPLA/Booster formulation. BCG booster of HspX/EsxS/PLUSCOM/MPLA was significantly induced higher levels of sIgA antibody secretion than Other BCG booster formulation, ISCOMATRIX/HspX/EsxS/MPLA (P <0.0001). Moreover, PLUSCOM containing formulations were able to induce higher sIgA responses than ISCOMATRIX containing formulation (Figure 7).

**Figure 7.**
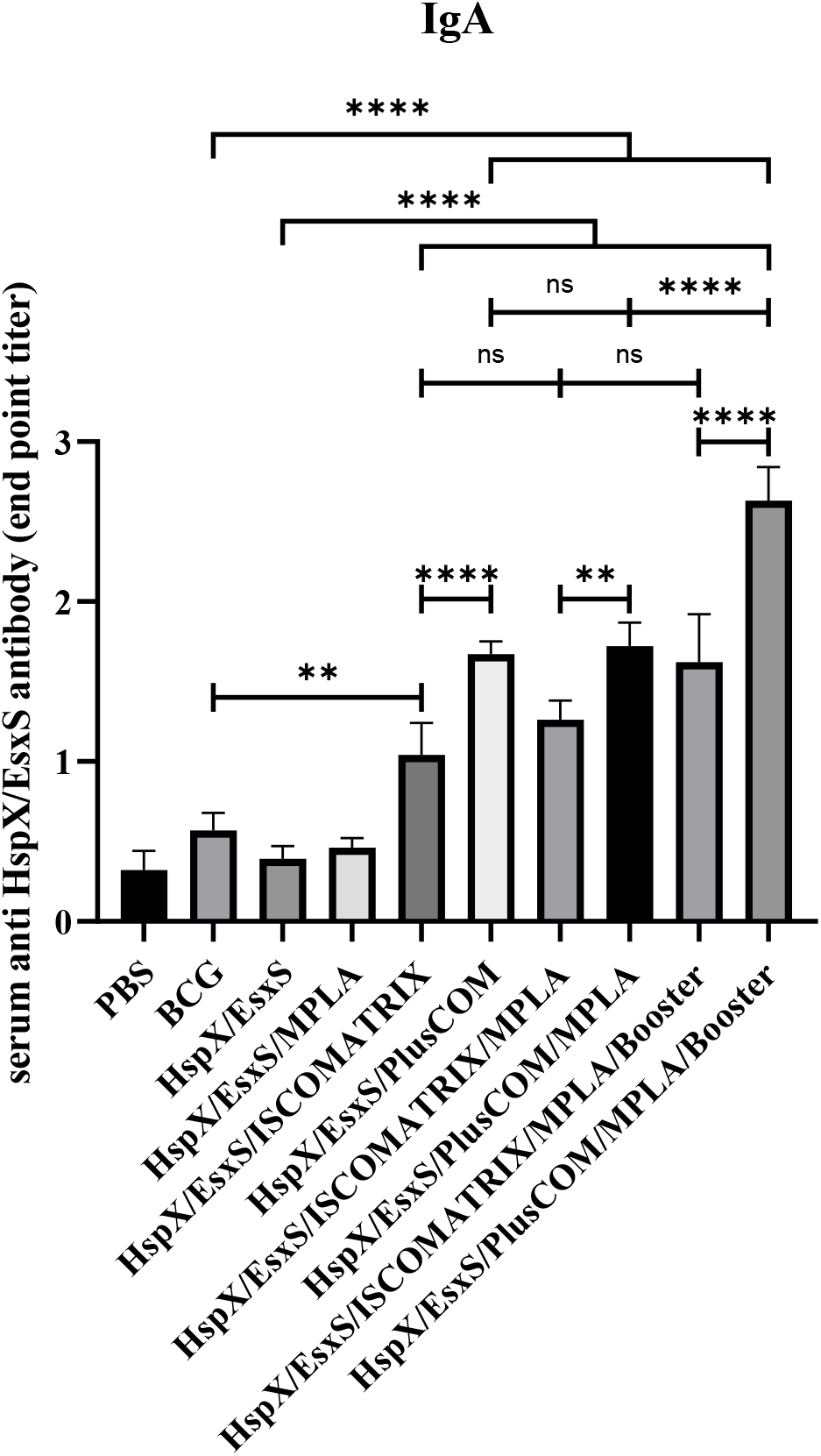
The level of anti-HspX/EsxS sIgA produced in nasal lavage of mice receiving different formulation.

## Discussion

After 1984 which Morein and colleagues for the first time were developed ISCOM-like structures, the results of several animal models and human clinical trials suggested that ISCOMATRIX-based vaccines are safe, well tolerated and immunogenic and able to induce strong humoral and cellular responses. The components of ISCOMATRIX adjuvant, i.e. saponin, cholesterol and phospholipid, form a cage-like structure (40-50 nm in diameter and about -20 mV in surface charge of particle) that facilitate antigen-presentation and antigen-delivery and also show immunomodulatory properties (25, 26). Efficacy of ISCOMATRIX adjuvant is currently under evaluation for cancer and some chronic infectious diseases such as hepatitis C virus and influenza, however, there is no study assessing ISCOMATRIX-based TB vaccines (27). In the current study, intranasal administration of ISCOMATRIX adjuvant in combination with HspX/EsxS antigen increased immune response especially the level of IFN-γ and IgG1, IgG2a and sIgA antibodies compared to alone antigen. A similar result was observed with the same formulation when administrated subcutaneously (20). It shows that ISCOMATRIX can boost immunogenicity of antigen which is a main weakness of subunit antigen vaccines. Other classic ISCOMs derivatives with a cage-like structure and positive surface charge is a cationic immune stimulating complex called PLUSCOM. The PLUSCOMs similar to ISCOMATRIXs can act as an immunoadjuvant and are able to induce T cell responses against an antigen, which is the most important human body response against TB infection (23, 28, 29). Positively charged PLUSCOM nano-adjuvant in combination with TB fused antigen was able to induce higher sIgA and IFN-γ responses than negatively charged ISCOMATRIX-antigen formulation after intranasal administration. Similar results were observed in subcutaneous route (20). One possible reason that is the positively charged PLUSCOM adjuvant strongly improves the particle-antigen uptake by the physiological surfaces such as mucosal surfaces as well as by the negatively charged immune cells particularly APCs and subsequent presentation to T cells (17, 28, 30). It is recommended that ISCOMATRIX adjuvant can be a good choice for using in the prophylactic and therapeutic vaccines. Prophylactic TB vaccine candidates are pre-exposure vaccines and similar to BCG can be administered after birth time. These types of TB vaccine candidates could be replaced with BCG or act as BCG booster (7, 25, 31). Our results revealed that ability of PLUSCOM/HspX/EsxS and ISCOMATRIX/HspX/EsxS formulations to elicit IFN-γ response were higher than BCG vaccine. These vaccine formulations cannot be replaced with BCG because the results were not statistically significant in some cases. Also, addition of MPLA adjuvant into ISCOMATRIX/HspX/EsxS and PLUSCOM/HspX/EsxS formulations was promoted the immune responses. The results were encouraging in intranasally vaccinated mice with formulations HspX/EsxS/ISCOMATRIX/MPLA, HspX/EsxS/PLUSCOM/MPLA and two BCG booster groups. Similar findings were obtained for the same groups when administrated via subcutaneous route (20).

## Conclusion

Taken together, our study suggested that ISCOMATRIX and PLUSCOM nano-adjuvants were able to boost HspX/EsxS immunogenicity and induced higher level of IFN-γ response and sIgA antibodies secretion compared to HspX/EsxS alone and addition of MPLA adjuvant promoted the immune responses. Furthermore, both HspX/EsxS/ISCOMATRIX/MPLA and HspX/EsxS/PLUSCOM/MPLA and their BCG booster formulation were able to induce IFN-γ response significantly higher than BCG group. These findings demonstrate that both nanoparticles in combination with MPLA can act as immunoadjuvant. However, further *in vivo* experiments are required to confirm the efficacy of these formulations as new TB vaccine or as BCG booster.

## Acknowledgment

This work was part of a PhD dissertation by Arshid Yousefi Avarvand and supported by a grant number 940965 from Vice Chancellor for Research, Mashhad University of Medical Sciences. The authors are grateful to personnel of Bu-Ali Research Institute, Mashhad University of Medical Sciences for technical assistance and valuable helps.

## Author contributions

Zahra Meshkat and Mohsen Tafaghodi conceived and designed research. Arshid Yousefi Avarvand conducted experiments. Arshid Yousefi Avarvand and Farzad Khademi analyzed data. Arshid Yousefi Avarvand wrote the manuscript. Ehsan Aryan, Mojtaba Sankian prepared the tables and figures. All authors read and approved the manuscript.

## Compliance with ethical standards

### Ethical statement

All applicable international, national, and institutional guidelines for the care and use of animals were followed.

